# Preclinical studies for plant-based oral enzyme replacement therapy (Oral-ERT) in Pompe disease knockout mice with transgenic tobacco seeds expressing human GAA (tobrhGAA)

**DOI:** 10.1101/2021.11.11.468227

**Authors:** Frank Martiniuk, Adra Mack, Justin Martiniuk, Shoreh Miller, Gregory O. Voronin, David Reimer, Nancy Rossi, Leslie Sheppard Bird, Sussan Saleh, Ruby Gupta, Mariel Nigro, Peter Meinke, Benedikt Schoser, Feng Wu, Angelo Kambitsis, John Arvanitopoulos, Elena Arvanitopoulos, Kam-Meng Tchou-Wong

**Author notes:** 551-999-06231, FM-whom reprints should be directed. shared equally in authorship and experiments (Adra Mack, Justin Martiniuk). (848) 445-8412-office (732) 233-2023-cell. Main Line: Direct Line.

## Abstract

Genetic deficiency of acid *α*-glucosidase (GAA) results in glycogen storage disease type II (GSDII) or Pompe disease (PD) encompassing at least four clinical subtypes of varying severity (infantile; childhood, juvenile and late onset). Our objective is to develop an innovative and affordable approach for enzyme replacement therapy (ERT) via oral administration (Oral-ERT) to maintain a sustained, therapeutic level of enzyme on a daily basis to improve efficacy of treatment and quality of life for people living with Pompe disease. A consensus at a 2019 US Acid Maltase Deficiency (AMDA) conference suggested that a multi-pronged approach including gene therapy, diet, exercise, etc. must be evaluated for a successful treatment of Pompe disease. Tobacco seeds contain the metabolic machinery that is more compatible with mammalian glycosylation-phosphorylation and processing. Previously, we have shown that a lysate from transgenic tobacco seeds expressing human GAA (tobrhGAA) was enzymatically active and can correct enzyme deficiency in cultured PD cells and in adult lymphocytes of Pompe patients and *in vivo* in disease-relevant tissues in GAA knockout (KO) mice when administered IP.

We have extended these pre-clinical studies in PD knockout (KO) mice with ground tobrhGAA seeds that supports proof-of-concept for Oral-ERT for future clinical trials. Briefly in GAA KO mice, Oral-ERT with ground tobrhGAA seeds showed significant reversal of fore-limb and hind-limb muscle weakness, increased motor coordination/balance/strength and mobility, improved spontaneous learning, increased GAA baseline activity in tissues, reduced glycogen in tissues and negible serum titers to GAA. Pharmacokinetics showed maximum serum GAA concentration (Cs) at 8-10 hr and peak urine excretion at 10-12 hr. The tobrhGAA was taken up in PD fibroblast, lymphoid and myoblast cells. Enzyme kinetics compared favorably or superior to placental hGAA, plus alglucosidase alfa or other rhGAAs for K_m_, V_max_, pH optima, thermal heat stability and IC_50_ for inhibitors. The tobrhGAA in seeds was extremely stable stored for 15 years at room temperature. NGS-genome sequencing of the tobrhGAA and wild-type plants and RNA expression profiles was performed and will be posted on our website. Thus, Oral-ERT with ground tobrhGAA seeds is an innovative approach that overcomes some of the challenges of alglucosidase alfa-ERT and provides a more effective, safe and significantly less expensive treatment.

## INTRODUCTION

Genetic deficiency of the lysosomal acid maltase or acid *α*-glucosidase (GAA) results in acid maltase deficiency (AMD) or Pompe disease (PD) encompassing at least four clinical subtypes of varying severity (infantile; childhood, juvenile and adult onset)(**1**). PD exhibits intracellular accumulation of glycogen in multiple tissues with skeletal muscle being the primary target, manifesting as myopathy and cardiomyopathy (**2-5**). The classic infantile form exhibits no enzyme activity, muscle weakness, feeding difficulties and hypertrophic cardiomyopathy, leading to death in the first year (**6-8**). The adult-onset form has partial enzyme deficiency, manifesting with slowly progressive muscle weakness leading to wheelchair/ventilator dependence and premature death from ventilator insufficiency (**9-11**). The incidence of people living with PD is estimated at 1:40,000 (**12**)(Orpha number-ORPHA365 at www.orpha.net). Recently, Park (**13**) re-calculated the carrier frequency (pCF) and predicted genetic prevalence (pGP) using general population databases based on the proportion of causative genotypes. Total pCF and pGP were 1.3% (1 in 77) and 1:23,232. The highest pGP was in the E. Asian population at 1:12,125, followed by Non-Finnish European (1:13,756), Ashkenazi Jewish (1:22,851), African/African-American (1:26,560), Latino/Admixed American (1:57,620), S. Asian (1:93,087) and Finnish (1:1,056,444). Unfortunately, the number of people in each population enrolled may not reflect the real-world population.

ERT with a recombinant human GAA (rhGAA) secreted by CHO cells (alglucosidase alfa/Myozyme/Lumizyme, Genzyme/Sanofi Corp.) infused every two weeks is the first approved therapy. Although it is efficient in rescuing cardiac abnormalities and extending the life span of infants, the response in skeletal muscle is variable (marketed dose of 20 mg/kg/every 2 weeks-iv) via the cation-independent mannose 6-phosphate receptor (**14-17**). In infants, muscle pathology and degree of glycogen deposition correlates with the severity of symptoms and the earlier ERT is introduced, the better chance of response (**14**). In adult-onset patients, mild improvements in motor and respiratory functions have been achieved which is unsatisfactory in the reversal of skeletal muscle pathology (**14-17**). In adult-onset patients, the ability for muscle to metabolize extra-lysosomal as well as intra-lysosomal glycogen is impaired. Lysosomal glycogen accumulation leads to multiple secondary abnormalities (autophagy, lipofuscin, mitochondria, trafficking and signaling) that may be amenable by long-term therapy. ERT usually begins when the patients are symptomatic and these secondary problems are already present contributing to inefficient delivery/uptake of alglucosidase alfa to muscle (**18**). The outcome of infantile patients is determined by many factors, among which are age, disease severity at ERT, genotype, genotype dependent CRIM status and the height of the antibody response. High and sustained antibody titers need to be prevented to achieve a good response to ERT, however titers varied substantially between patients and does not strictly correlate with the patients’ CRIM status. Only infantile patients can be either CRIM-negative or CRIM-positive. Since all juvenile and adult patients make at least some amount of residual GAA, they are always CRIM-positive. About 40% of infantile patients are CRIM-negative while the remaining 60% are CRIM-positive. The immune response may be minimized by early start of ERT and pretreatment by an immune tolerance induction regime using e.g. a combination of rituximab, methotrexate, bortezomib and IV immunoglobulins. Up to 20% of adult patients develop high titers on ERT, 40% intermediate and 40% none or low titers (**19-40**). In adults, antibody formation does not interfere with rhGAA efficacy in the majority of patients, however may be associated with infusion-associated reactions (IARs) and may be attenuated by the IVS1/delex18 (c.2481+102_2646+31del, expression of a truncated protein) GAA genotype (**41**). Some patients experienced IARs due to bronchial spasm with flushing and pulmonary artery blockage during infusion. Several studies suggest that unchanged endogenous GAA protein expression lowers the chance of forming antibodies to recombinant proteins. Gutschmidt et al. (**42**) found that ERT in late onset patients’ clinical outcomes, particularly lung function, muscle strength and walking capability tend to deteriorate over time, indicating that additional efforts must be made to improve ERT effectiveness and supplement therapies developed. Korlimarla et al (**43**) used a standardized behavior checklists as screening tools for the early identification and treatment of behavior, emotional and social concerns in children with PD. Although alglucosidase alfa has been a wonderful first step in treating people with PD, it has revealed subtle aspects that must be dealt with for successful treatment (**18,43-49**). ERT has unmasked previously unrecognized clinical manifestations include tracheo-bronchomalacia, vascular aneurysms and GI discomfort that impacts smooth muscle. Persistent smooth muscle pathology has a substantial impact on quality of life and leads to life threatening complications. In addition to airway smooth muscle weakness, vascular deterioration, GI discomfort and loss of genitourinary control have been observed. Cerebral and aortic aneurysms have caused microhemorrhages leading to symptoms ranging from headaches and numbness to coma and death (**50-56**). PD results in subsequent pathology in smooth muscle cells may lead to life-threatening complications if not properly treated (**57**). In GAA KO mice, there is increased glycogen in smooth muscle cells of the aorta, trachea, esophagus, stomach and bladder plus increases in lysosome membrane protein (LAMP1) and autophagosome membrane protein (LC3). More importantly, lifelong treatment ($250-500k/yr/adult) can be prohibitively expensive resulting in the reluctance of insurance companies to reimburse costs for adult patients (**58**) underlining the demand for more economical production and novel delivery strategies for treatment. A consensus at a 2019 US Acid Maltase Deficiency Association conference in San Antonio, TX, suggested that a multi-pronged approach including gene therapy, diet, exercise, etc. must be evaluated for a successful treatment of PD.

The technological platform involving the accumulation of recombinant proteins in seeds warrants better availability of the products and allows long-term storage of the biomass for processing (**59-61**). Transgenic plants, seeds and cultured plant cells are potentially one of the most economical systems for large-scale production of recombinant enzymes for pharmaceutical uses (**61-69**). Seeds are particularly attractive due to their high rates of protein synthesis and their ability to remain viable in the mature-dry state as a stable repository (**68-81**). Over one-third of approved pharmaceutical proteins are glycoproteins (**73-83**) and even minor differences in N-glycan structures can change the distribution, activity or longevity of recombinant proteins when compared with their native counterparts, altering their efficacy as therapeutics (**60,61,84-87**). Protalix Biotherapeutics and Shaaltiel et al. (**87,88**) produced a glucocerebrosidase for ERT of Gaucher disease using a plant cell system and an Exploratory, Open-label Study to Evaluate the Safety of PRX-112 and PK of oral prGCD in patients receiving 250 ml of a re-suspended carrot cells. Administration of PRX-112 using carrot cells as carrier vehicle, may overcome absorption, degradation and uptake in the small intestine and can be found in the blood stream in an active form (**89-91**).

Previously, we expressed the human *GAA* cDNA in chloroplasts, callus and leaves of transgenic tomato and tobacco plants but was not enzymatically active. The *hGAA* was expressed in tobacco seeds (tobrhGAA) by targeting to the ER using the signal peptide sequence/promoter of soybean β-conglycin (**92**). The tobrhGAA from transgenic plant seeds was enzymatically active, taken up by PD fibroblasts and WBCs to reverse the enzyme defect and binds to the affinity matrix Sephadex G100 and eluted by maltose. A crude lysate of transgenic seeds administered intraperitoneally in GAA KO mice increased GAA activity to 10-20% of normal in tissues, most notably heart, skeletal muscle and diaphragm (**92-97**).

In this report, we have extended these pre-clinical studies in PD KO mice with ground tobrhGAA seeds that supports proof-of-concept for Oral-ERT for people living with PD for future clinical trials. Briefly in GAA KO mice, Oral-ERT with various preparations of tobrhGAA showed significant reversal of fore-limb and hind-limb muscle weakness, increased motor coordination/balance/strength and mobility, improved spontaneous learning, increased GAA baseline activity in tissues, reduced glycogen in tissues and negligible serum titers to GAA. Pharmacokinetics showed maximum serum GAA concentration (Cs) at 8-10 hr and peak urine excretion at 10-12 hr. The tobrhGAA was taken up in fibroblast and lymphoid cells from infantile, juvenile and adult onset patients comparable to human placenta GAA and an rhGAA. Exposure of tobrhGAA to a human PD myoblast cell line increased GAA to 24-35% of normal. Enzyme kinetics for tobrhGAA vs placental hGAA, plus alglucosidase alfa or other rhGAAs for K_m_, V_max_, pH optima, thermal heat stability and IC_50_ for inhibitors and our tobrhGAA is comparable to placental GAA and superior to alglucosidase alfa and other rhGAAs. The tobrhGAA in seeds stored for 15 years at room temperature showed less than 15% loss in GAA activity indicating extreme stability. NGS-genome sequencing of the tobrhGAA and wild-type plants and RNA/proteome expression profiles will be posted on our website (under construction). Oral-ERT with tobrhGAA is an innovative approach that overcomes some of the challenges of alglucosidase alfa-ERT and provides a more effective, safe and less expensive treatment. This protocol will be the second oral ERT plant-made pharmaceutical (PMPs)(**84,89-91,96**).

## MATERIAL and METHODS

### Hydroponic Growth of Tobacco Plants

Transgenic tobacco plant #3 expressing human GAA in seeds are grown indoors using a hydroponic system (Active Aqua Grow Flow Kit, Hydrobuilder, Inc.) with deionized water (eliminating water/soil contaminants). Seed-pods are harvested and dried in freeze-dryer followed by separation from the husk by passage through a standard food mesh strainer. To eliminate environmental contamination, seeds were ground for 10 minutes in a porcelain pestle-mortar until a fine powder and placed in a UV germicidal incubator for 30 minutes to sterilize. Viability of 0.1 grams (∼1,000 whole seeds) are grown in a 150 mm petri dish on paper towels saturated with water for 7-8 days at room temperature.

### Simulated Stomach and Small Intestinal Environments

To mimic the stomach and small intestine (SI) environment, we exposed the 100 mg of whole or milled tobrhGAA seeds to physiologic conditions/times to pepsin (stomach) followed by trypsin/chymotrypsin (small intestine)(**88,97**). Samples were added to 300 μg/ml pepsin at pH 4.0 for 60 min., adjusted pH to 6.5 and then trypsin at 800 μg/ml and chymotrypsin at 700 μg/ml for 60 min at 37°C followed by GAA assay.

### Enzyme Assay

Ground seeds or tissues were resuspended in 10 mM sodium phosphate, pH 7.5, frozen and thawed 3x and centrifuged at 13,000g for 10 min. The supernatants were assayed for GAA using the artificial substrate 4-methylumbelliferyl-*α*-D-glucoside (4-MU-Glc, 1 mg/ml) at pH 4.0 (0.5 M sodium acetate) and as an internal control, neutral alpha glucosidase (NAG) at pH 7.5 (0.5 M sodium phosphate) for 18 hours. Fluorescence was determined in a fluorometer (excitation-360_nm_ and emission-460_nm_)(Sequoia-Turner) as previously described (**98**).

### Biochemical/Enzyme Kinetic Analyses

We compared a lysate of tobrhGAA seeds to a rhGAA (R&D Systems) and mature placental human GAA for specific activity using 4-MU-Glc at pH 4.0, maltose and glycogen, pH optima, inhibitors (acarbose, castanospermine, deoxynojirimycin, miglitol, voglibose) and heat stability (**95,99-109**) using standard enzyme kinetic methods.

### Uptake of tobrhGAA by PD Human Myoblast, Fibroblast and Lymphoid Cell Lines

Human lymphoid (GM6314, GM13793, GM14450) or fibroblast (GM4912, GM1935, GM3329) cell lines from infantile or adult PD were maintained in 15% fetal bovine serum, RPMI 1640 or DMEM supplemented with glutamine, penicillin and streptomycin at 37°C-5% CO_2_. Cells were plated at 0.3-0.4 × 10^6^/6-well in 1.5 ml media-24 hr before addition of varying amounts of tobrhGAA or others GAA formulations (**93**). Cells were harvested after various hours of exposure, washed with PBS, lysed by addition of 0.5 ml of 0.01 M sodium phosphate pH 7.5, frozen and thawed 3x, spun for 5 minutes to clarify and assayed for human GAA and NAG as described above. A human PD myoblast cell line (homozygous for the IVS1 c.-32–13t>g)(**110**) and normal skeletal muscle cells were plated at 0.3 to 0.4 × 10^6^ in a 6-well plate with PromoCell skeletal muscle growth media. Cells were exposed a lysate of tobrhGAA for 48 hr and assayed. Mock treated GAA and normal myoblast cells were controls plus cells treated with equivalent amounts of a rhGAA (R&D Systems #8329-GH-025). Simultaneously, we measured cell proliferation with the Cayman MTT cell proliferation assay (#10009365).

### Short-term Studies GAA KO Mice

We utilized the GAA KO mouse with the exon 6^neo^ disruption (**111**), wild-type BALB/c or 129/J or GAA KO mice mock-treated with PBS. GAA KO mice (∼4-6 months old) were given oral administrations every other day to day 7 a lysate from 300 mg (∼75 μg tobrhGAA) of transgenic seeds mixed 3:1 with apple juice as a safe, non-invasive oral administration technique (**111,112**) and grip-strength measured by a grip-strength meter (GSM)(Columbus Inst. OH). At day 7, mice were sacrificed and tissues assayed for GAA and NAG and compared to wild-type mice and mock (PBS) treated GAA KO mice.

### Long-term Treatment by Oral Gavage in GAA KO Mice

We treated two groups (age matched) of GAA KO mice (exon 6^neo^)(n=3) three times weekly with either a 1x or 3x dose (containing either 25 or 75 μg tobrhGAA per dose) mixed 3:1 with apple juice as a safe, non-invasive oral administration technique (**101**) and measured vertical hang-time as performed by Raben et al. (**111**).

### Oral-ERT in GAA KO Mice

GAA KO mice were fed either 50 mg or 150 mg tobrhGAA ground seeds-daily mixed with peanut butter in petri dishes (12:12 L/D photoperiod)(**92**)(age ∼5 months)(B6,129-Gaa^tm1Rabn^/J mice, **110**). Tissues, urine, blood slides, weight and serum were collected. Mice were tested every 3 weeks for: motor activity by running wheel (RW-Mini-Mitter Co., Inc., OR); fore-limb muscle strength by grip-strength meter (GSM)(Columbus Inst. OH), motor coordination/balance with a Rotarod (AJATPA Expert Industries, India), open-field mobility by a 5 minute video for distance traveled and spontaneous alternative learning with a T-maze (Stoelting, IL).

### ELISA for Antibodies to tobrhGAA

In Falcon plates #3912, add Sephadex G100 purified human placental GAA (**113**) at 1 μg/well in 100 μl PBS at 4°C overnight, wash 3x with PBS, block with 200 μl PBS-1g% BSA-0.05% Tween-20 for 60 min. or longer at RT. Add 1 μl serum in 100 ul PBS-overnight at 4°C. Wash 3x PBS, 1x PBS-0.05% Tween-20, 2X PBS. Add 100 μl goat anti-mouse IgG-HRP (BioRad #172-1101) at 1:500 in PBS/BSA/Tween-20 at 4°C overnight. Wash 3x PBS, 1x PBS-0.05% Tween-20, 2x PBS. Stain with 100 μl of 3,3’,5,5’–tetramethylbenzidine (Sigma #T5525) in 9 ml of 0.05 M phosphate-citrate buffer, pH 5.0 (Sigma #P4922) with 20 μl of 30% H_2_O_2_ for 30-60 minutes at room temperature. Reactions are stopped with 50 μl 1 N HCl and read at A_450_nm.

### Endotoxin Determination

Endotoxin in various amounts of a lysate of tobrhGAA seeds was determined using the Pierce LAL Chromogenic Endotoxin Quantitation Kit (#88282) according to manufacturers instructions (**114**).

### Bradford protein assay

150 ul reagent (BioRad #500-0006) diluted 1->5 with water in a 96 well microtiter plate. Standards were serial dilutions of BSA from 1,000 μg/ml to 2 μg/ml and 1 to 25 μl samples. Samples were read at A_595_nm before 60 min.

### Assay for Glycogen Content

In a 96-well microtiter plate, add 25 μl sample, 25 μl 0.05 M sodium phosphate pH 6.5, ±0.5 μl amyloglucosidase (Sigma #A-1602), placed overnight at 37°C (**115**). Glycogen standards (rabbit liver, 2 mg/ml) and D-glucose standards were at 400 μM, 200 μM and 100 μM. Add 20 μl of Eton Bioscience glucose assay solution (#1200031002) and incubate for 30 minutes at 37**°**C. The reaction was stopped with 25 μl of 0.5 M acetic acid and read at A_490_nm.

### Ames Mutagenicity Test

on *E*.*coli* DH5*α* and DH5*α*/pUC19 plated on NZ media with ampicillin and various amounts of tobrhGAA lysate and colony forming units counted (**114,116-119**).

### CBC differentials

CBC differentials were perform on blood smears on slides with Giemsa-Wright staining.

### Extracton of DNA and RNA from Seeds for NGS and RNA Transcriptome profile

DNA and total RNA was extracted from wild-type (*Nicotiana tobacum L. cv. xanthi*) and tobrhGAA seeds by kits (Thermo Scientific GeneJET Plant RNA purification kit-K0801; GeneJET Plant DNA purification kit-K0791). The NGS-genome sequence and RNA expression profiles was performed at BGI.com and global proteomic profiling by UPLC-MS/MS at bioproximity.com.

### Nicotine Levels in Leaves and Seeds

The levels of nicotine in tobrhGAA#3 seeds and leaves was measured by gas chromatography-mass spectroscopy (GC/MS) at Avogado nAnalytical, LLC, Salem, NH.

### Statistical Analysis

Student t-test for probability associated with a population and standard deviation based upon the population were determined using Microsoft Excel software and considered significant at p = ≤0.05.

## RESULTS

We have extended our original studies in the production of a functional human acid maltase in tobacco seeds: biochemical analysis, uptake by human PD cells, and *in vivo* studies in GAA knockout mice (**92**) that is organized into three categories-**A**-Miscellaneous studies, **B**-*in vitro* studies and **C**-*in vivo* studies which evolved from the use of whole seeds or seed lysates to the final ground seeds.

### A- Miscellaneous Studies

#### A.1 Hydroponic growth of tobacco plants and processing of seeds

Transgenic tobacco plant #3 expressing human GAA in seeds are grown indoors using a hydroponic system with deionized water (eliminating water/soil contaminants). Seed-pods are harvested and dried in freeze-dryer followed by separation from husk through a standard food mesh strainer. To eliminate environmental contamination, ground seeds were tested for lack of viability and germination. Seeds are grounded for 10 minutes in a porcelain pestle-mortar until a fine powder followed by UV sterilization. Approximately, 0.1 gram of ground/sterilized seeds (∼1,000 whole seeds) and untreated transgenic seeds are germinated (positive control) by evenly distributed in a petri dish on paper towels saturated with water and observed for 7-8 days at room temperature. Viable seeds will germinate in a week (**Fig. 1**). No germination was observed in processed seeds.

**Figure 1.**
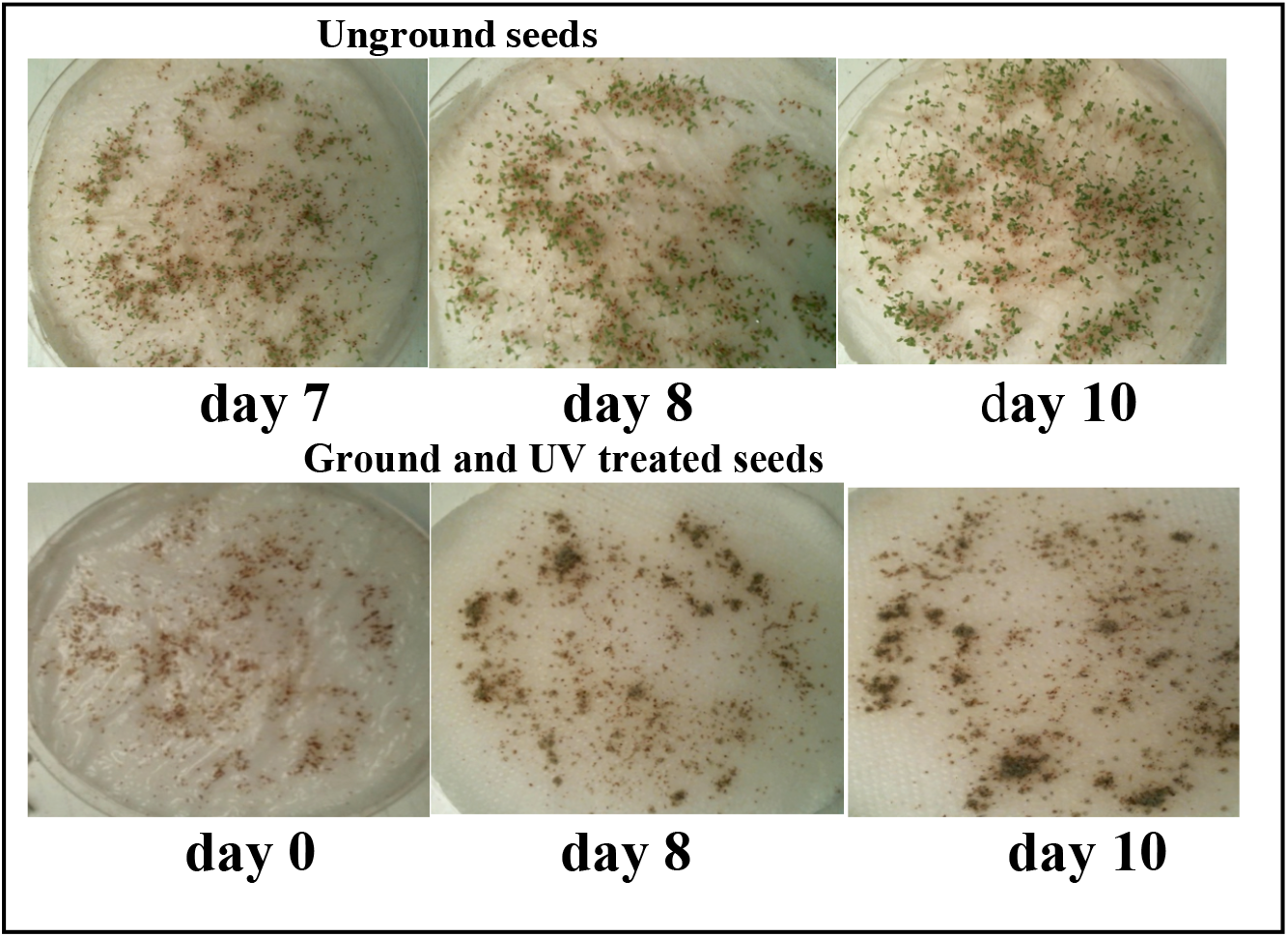
Hydroponic growth of tobacco plants and processing of seeds. Transgenic tobacco plant #3 expressing human GAA in seeds are grown indoors using a hydroponic system with deionized water (eliminating water/soil contaminants). Seed-pods are harvested and dried in freeze-dryer followed by separation from husk through a standard food mesh strainer. To eliminate environmental contamination, ground seeds were tested for lack of viability and germination. Seeds are grounded for 10 minutes in a porcelain pestle-mortar until a fine powder and placed in a UV germicidal incubator for 30 minutes to sterilize. Approximately, 0.1 gram of ground/sterilized seeds (∼1,000 whole seeds) and untreated transgenic seeds are germinated (positive control) by evenly distributed in a petri dish on paper towels saturated with water and observed for 7-8 days at room temperature. Viable seeds will germinate in a week. No germination was observed in processed seeds

#### A.2 Nicotine levels in leaves and seeds

was measured by GC/MS (Avogado nAnalytical, LLC, Salem, NH) and contained <5ng/dry gram of tobrhGAA seeds or leaves and generated a GC/MS spectral profile for future comparisons.

#### A.3 Long-term stability

We compared tobrhGAA in seeds stored for 9 and 15 years at room temperature vs. freshly harvested seeds. We found that there was less than 15% loss in GAA activity between the seeds indicating extreme stability (old = 0.20 μg tobrhGAA/g seeds vs. fresh-0.25 μg tobrhGAA/g seeds) for both age groups.

#### A.4 WGS, RNA and proteomic profiles

The WGS genome sequence and RNA expression profiles of the tobrhGAA#3 and wild-type plants were performed at BGI.com and global proteomic profiling by UPLC-MS/MS at bioproximity.com. Proteomic profile showed 285 proteins expressed in wild-type and tobrhGAA #3 seeds. Besides the hGAA, ∼20 proteins were found differentially expressed in tobrhGAA#3 seeds. Once the analyses and comparisons are completed, the data will be shared and posted on our website (under construction).

#### A.5 Endotoxin levels

in an extract of tobrhGAA #3 seeds with an LAL endotoxin kit was less than 0.25 EU/ml endotoxin or ∼25 pg/ml, values much lower than the estimate of 300 mg/kg for oral endotoxin toxicity in mammals and humans (**119-121**). We also found no viable anerobic or aerobic bacteria.

#### A.6 Affect of stomach and small intestinal environments

To mimic the stomach and small intestine (SI) environments, we exposed the tobrhGAA to physiologic conditions and times to pepsin (stomach) and trypsin/chymotrypsin (small intestine)(**88,97,122,123**). Lysosomal GAA is stable at low pH. We exposed a lysate from tobrhGAA #3 seeds to 300 μg/ml pepsin at pH 4.0 followed by trypsin at 800 μg/ml-pH 6.5/chymotrypsin at 700 μg/ml-pH 6.5 for 60 min at 37°C. None of the enzymes had any affect on tobrhGAA activity, thus demonstrating that the conditions in the digestive tract probably does not affect tobrhGAA. We repeated conditions with 100 mg of whole or milled seeds. Interestingly, the amount of tobrhGAA in whole and milled seeds (600 μg/g seeds) was more than double physical lysing suggesting that the acid environment in the stomach more thoroughly disrupts and releases tobrhGAA from the whole or milled seed. These data are extremely important for future production, manufacturing, costs and availability via a pill containing whole or milled seeds.

#### A.7 Ames mutagenicity test

was performed on *E. coli* DH5*α* and DH5*α*/pUC19 with various amounts of tobrhGAA seed lysate. We found no additional bacterial or plasmid revertant CFUs generated with a tobrhGAA lysate by exposure to >10^4^ bacteria (**114,118,119,123,124**).

### B- *in vitro* Studies

#### *B*.*1 in vitro* studies in PD fibroblast and lymphoid cell lines

We evaluated tobrhGAA uptake in fibroblast and lymphoid cells from infantile, juvenile and adult onset patients and compared to human placenta GAA and an rhGAA (R&D Systems). Various equivalent concentrations and time-points (2, 24 and 48 hr) showed similar uptake and increases in GAA (mean±SEM)(**Graphs 1 and 2**).

#### B.2 Uptake of tobrhGAA in a human PD myoblast cell line

A human PD myoblast cell line (homozygous for the IVS1 c.-32–13t>g)(**110**) was exposed to equivalent amounts of a lysate of tobrhGAA seeds for 48 hr and assayed. Mock treated GAA and normal skeletal muscle cells were controls plus cells treated with rhGAA. We found that tobrhGAA increased GAA to 24-35% of normal (mean±SD)(**Graph 3**). Simultaneously, we measured cell proliferation with the Cayman MTT assay and found that both tobrhGAA doses slightly reduced cell growth by <10%.

#### B.3 Biochemical/enzyme kinetic analyses

We compared the enzyme kinetics for a lysate of tobrhGAA vs placental human placental GAA and an rhGAA (R&D Systems), plus limited published data for alglucosidase alfa or other rhGAAs (**101-109**) for K_m_, V_max_, pH optima, thermal heat stability and IC_50_ for inhibitors (acarbose, castanospermine, deoxynojirimycin, miglitol, voglibose). Data showed that showed our tobrhGAA is comparable to placental GAA and superior to alglucosidase alfa and other rhGAAs (**Table 1**)(**95,99-109,113,125,126**).

**Table 1.**
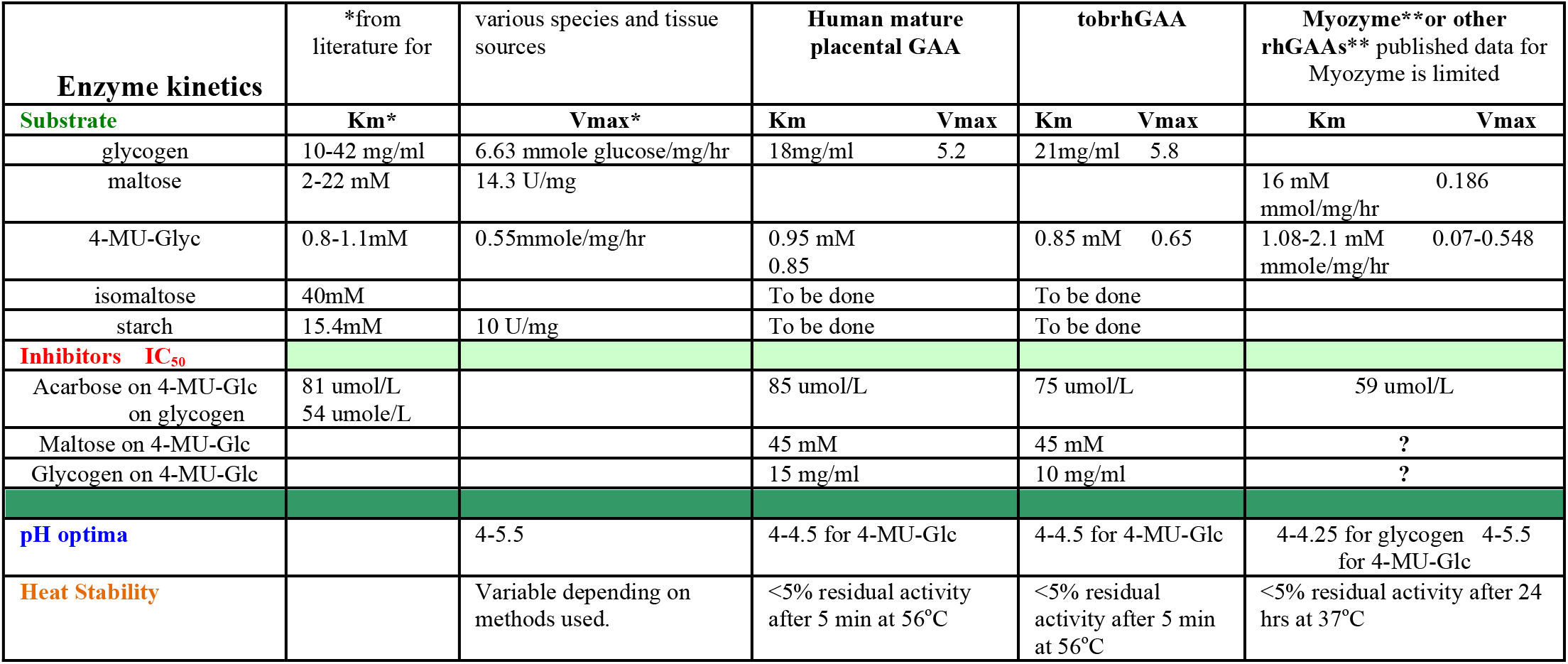

### C- *in vivo* Studies

#### C.1 Short-term treatment

Fore-limb grip strength was measured by a grip strength meter in GAA KO mice (exon 6^neo^)(**111**) after three oral administrations every other day to day 7 with a lysate (1x dose containing 75 μg tobrhGAA) of tobrhGAA seeds. Wild-type mice (n=3) averaged 245±21 lbs (SEM) grip at release. Mock treated GAA KO mice (n=3) averaged 92±3 lbs grip at release (3-5 attempts/mouse) and treated GAA KO mice (n=3) averaged 105±3 lbs grip at release (p = ≤0.024 treated vs. mock treated) with a 14% improvement in strength. At day 7, mice were sacrificed and tissues were assayed for GAA/NAG and compared to mock-treated GAA KO and wild-type mice. Importantly, we found increases in GAA activity in tissues from 8 to 23% of wild-type mice (heart, skeletal muscle, liver and diaphragm) from treated GAA KO mice (**Table 2**).

**Table 2.** GAA/NAG (mean ± SD) assay of mouse tissues after administration of tobrhGAA for 3x over 7 days via oral gavage.

|  | Hind-limb tibialis anterior<br>Skeletal Muscle | Heart | Diaphragm | Liver |
| --- | --- | --- | --- | --- |
| Treated GAA KO - Oral tobrhGAA | 0.12 ± 0.04 | 0.10 ± 0.06 | 0.13 ± 0.09 | 0.19 ± 0.02 |
| Mock GAA KO- PBS | 0.043 ± 0.006 | 0.06 ± 0.01 | 0.07 ± 0.03 | 0.05 ± 0.009 |
| Wild-type-Balb/c | 1.50 ± 0.2 | 0.43 ± 0.19 | 0.77 ± 0.15 | 1.00 ± 0.13 |
| % of WT | 8% | 23% | 17% | 19% |

#### C.2 Long-term (203 days) treatment

We treated two groups of GAA KO mice (**111**)(n=3)-3x weekly with either a 1x or 3x dose (25 or 75 μg tobrhGAA/dose) orally (**101**) and measured vertical hang-time over 203 days. The treatment was well tolerated as mice gained weight during the treatment. Both treatment groups showed a steady significant improvement in vertical hang-time vs mock treated GAA KO mice. Interestingly, mice in the 1x group gradually increased hang-time equal to the 3x dose group, suggesting that the lower dose can be effective in long-term treatment. Both treatment groups showed 25-35% vs. mock treated wild-type mice at day 82, but by day 203, both groups were almost equal (89-95%) to mock treated wild-type mice (**Fig. 2**).

**Figure 2.**
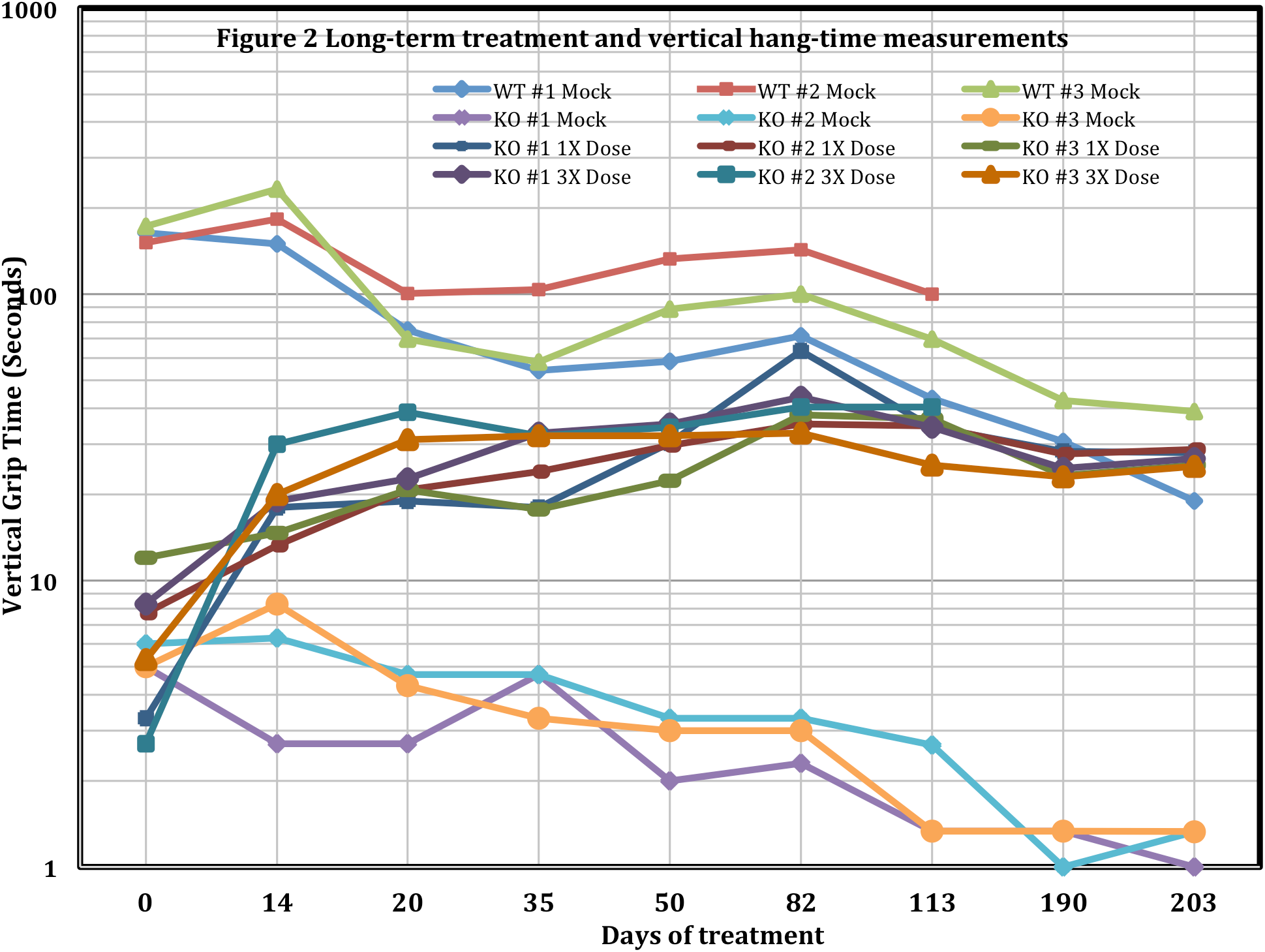
Long-term treatment and vertical hang-time measurements. We treated 2 groups of GAA KO mice (exon 6^neo^)(n=3)-3x weekly with either a 1x or 3x dose (25 or 75 μg tobrhGAA/dose) orally and measured vertical hang-time over 203 days. Both treatment groups showed a steady significant improvement in vertical hang-time vs. mock treated KO mice. Interestingly, mice in the 1x group gradually increased hang-time equal to the 3x dose group, suggesting that the lower dose can be effective in long-term treatment. Both treatment groups showed 25-35% vs. mock treated wild-type mice at day 82, but by day 203, both groups were almost equal (89-95%) to mock treated wild-type mice.

##### Assay of tissues at 203 days treatment

We found increases in GAA activity in tissues (heart, hind-limb skeletal muscle, kidney, liver and diaphragm) from GAA KO mice treated with tobrhGAA by oral gavage, thereby supporting Oral-ERT with tobrhGAA. The % of wild-type ranged from 13 to 38% depending upon tissue (**Table 3**).

**TABLE 3.**
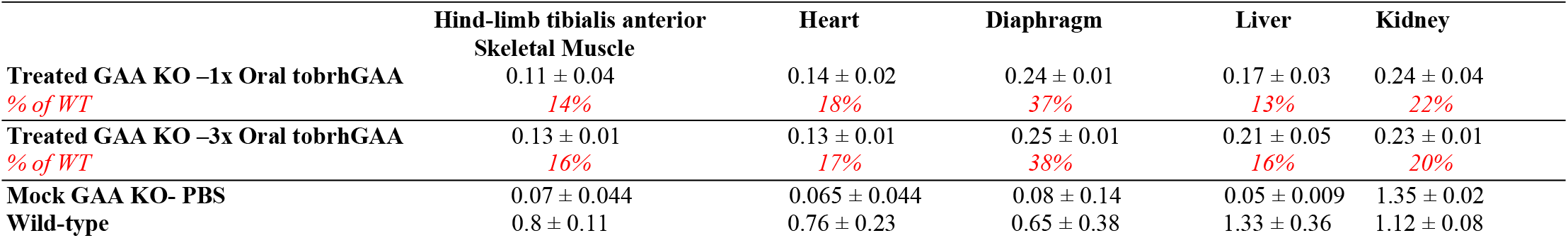
GAA/NAG (mean ± SD) assay of mouse tissues after administration of tobrhGAA via oral gavage (Oral-ERT) at day 203.

|  | Hind-limb tibialis anterior<br>Skeletal Muscle | Heart | Diaphragm | Liver | Kidney |
| --- | --- | --- | --- | --- | --- |
| Treated GAA KO -1x Oral tobrhGAA | 0.11 ± 0.04 | 0.14 ± 0.02 | 0.24 ± 0.01 | 0.17 ± 0.03 | 0.24 ± 0.04 |
| % of WT | 14% | 18% | 37% | 13% | 22% |
| Treated GAA KO -3x Oral tobrhGAA | 0.13 ± 0.01 | 0.13 ± 0.01 | 0.25 ± 0.01 | 0.21 ± 0.05 | 0.23 ± 0.01 |
| % of WT | 16% | 17% | 38% | 16% | 20% |
| Mock GAA KO- PBS | 0.07 ± 0.044 | 0.065 ± 0.044 | 0.08 ± 0.14 | 0.05 ± 0.009 | 1.35 ± 0.02 |
| Wild-type | 0.8 ± 0.11 | 0.76 ± 0.23 | 0.65 ± 0.38 | 1.33 ± 0.36 | 1.12 ± 0.08 |

##### Glycogen reduction in tissues

We measured glycogen levels in tissues (**110**) after 203 days treatment and found it ranged from 18 to 48% reduction depending upon tissue (**Table 4**).

**TABLE 4.** % glycogen reduction (mean) in mouse tissues after administration of tobrhGAA at day 203 (Oral-ERT).

|  | Hind-limb tibialis anterior<br>Skeletal Muscle | Heart | Diaphragm | Liver | Kidney |
| --- | --- | --- | --- | --- | --- |
| Treated GAA KO -1x Oral tobrhGAA | 39 | 32 | 48 | 18 | 28 |
| Treated GAA KO -3x Oral tobrhGAA | 43 | 18 | 24 | 25 | 29 |

##### Immune response

Serum antibody titer to oral tobrhGAA at 203 days was undetectable at 4 weeks, barely detected at a 1:10 at 8 weeks for both 1x and 3x doses and detected at 1:20, thus demonstrating low immune response (**102**). We simultaneously generated mouse antibody to human placental GAA by s.c. injection of 15 ug over 3 months and found the titer to be 1:1500. Thus, oral tobrhGAA of similar quantity of antigen did not generate a high serum immunoreaction.

##### Taken up by a distant tissue

We determined if the tobrhGAA was taken up by a distant tissue-tail without sacrificing the mice and assayed for GAA at 45 and 203 days. Wild-type mice had GAA activity of 0.46±0.14, mock treated GAA KO had 0.064±0.021 and treated GAA KO had 0.10 to 0.2±0.05 for 1x or 3x dose (p=≤0.006 treated vs mock treated)(**Graph 4**).

#### C.3 Treatment with whole seeds and running wheel activity

GAA KO mice (3 females-∼4 months age) were fed twice daily with 50 mg whole tobrhGAA seeds and running wheel (RW) activity monitored for 21 days, followed by 50 mg-1x daily for 14 days and then 50 mg every 2 days (**Fig. 3**). Historical levels for wild-type and GAA KO mice and GAA KO mice feed 50 mg of wild-type tobacco seeds are indicted. GAA KO mice treated 2x daily with 50 mg tobrhGAA seeds showed increased running wheel activity to 60% of wild-type by day 24. At day 25, mice were feed 50 mg-1x daily, resulted in a reduction of RW to 40% of wild-type and at day 41, mice were feed 50 mg every other day-reduced activity to mock treated levels.

**Figure 3.**
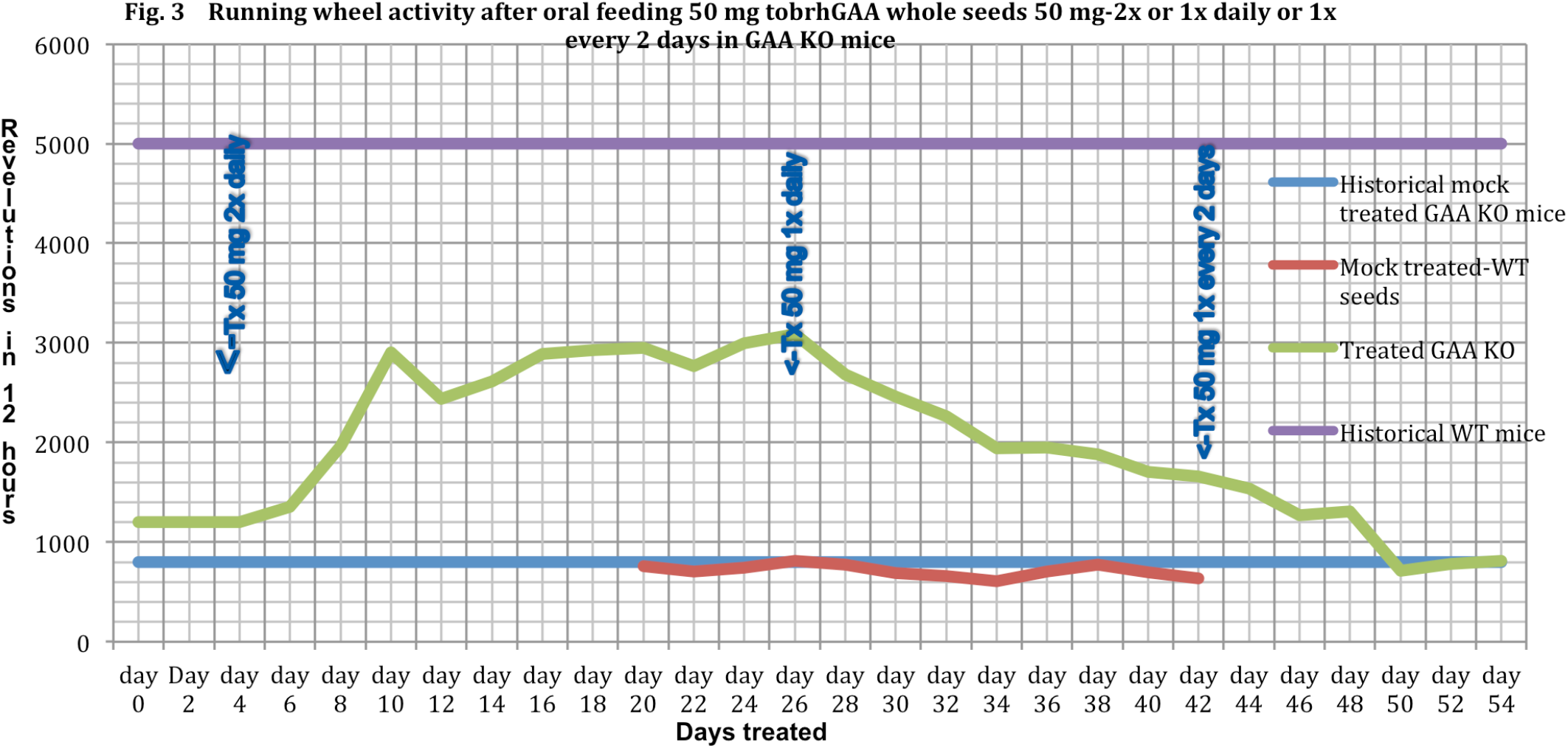
Treatment with whole seeds and running wheel activity. GAA KO mice (3 females-∼4 months old) were fed twice daily with 50 mg whole tobrhGAA seeds and running wheel activity monitored for 21 days, followed by 50 mg-1x daily for 14 days and then 50 mg every 2 days. Historical levels for wild-type and GAA KO mice and GAA KO mice feed 50 mg of wild-type tobacco seeds are indicted. GAA KO mice treated 2x daily with 50 mg tobrhGAA seeds showed increased running wheel activity to 60% of wild-type by day 24. At day 25, mice were feed 50 mg-1x daily, resulted in a reduction of RW to 40% of wild-type by day 40 and at day 41, mice were feed 50 mg every other day-reduced activity to mock treated levels.

##### C.4 Assessment of spontaneous alternation in wild-type and GAA KO mice

Spontaneous alternation is used to assess the cognitive ability of rodents to choose one of the 2 goal arms of the T-maze. The advantage of a free choice procedure is that hippocampal or lesioned animals often develop a side preference and scores below 50%. Controls generally achieve at 60-80% correct alternation. We assessed spontaneous alternative learning for cognitive ability in the T-maze in both male and female GAA KO mice and wild-type-129/C57 mice from 2-9 months of age. We found that deficiency in spontaneous learning appeared by 2-3 months in male and 3-4 months in female GAA KO mice (**Graph 5**). All conditions were significant (p=≤0.05).

##### C5. Oral-ERT with tobrhGAA in GAA KO mice

GAA KO mice were administered 50 mg or 150 mg tobrhGAA ground sterile seeds-daily (age 5 months)(B6,129-Gaa^tm1Rabn^/J mice, **111**). Tissues, urine, blood slides, weight and serum were collected. Mice were tested every 3 weeks for: motor activity by running wheel (RW); fore-limb muscle strength by grip-strength meter (GSM), motor coordination/balance with a rotarod, open-field mobility by 5 min video/distance traveled and spontaneous learning with a T-maze. Reversal of muscle weakness showed that young mice responded faster than older mice and older mice required higher levels of tobrhGAA. Both age groups and doses showed improvement in spontaneous alternation by T-maze after 2-3 weeks (**Table 5**). CBC differentials of treated GAA KO animals were identical to pre-treatment GAA KO and wild-type-129/C57 mice.

**Table 5.**

| Table 5 | RW | GSM | Rotarod | T-maze | Open-field mobility | CBC diff. |
| --- | --- | --- | --- | --- | --- | --- |
| Mean±SD | rev/12 hrs | lbs at release | Time (sec) | % correct alternation | cm/min ±SD |  |
| Pre-Tx n = 7-37 M/F | 404±128 | 0.80±59 | 25±49 | 32±25 | 182±79 |  |
| GAA KO M/F-50mg n = 4 |  |  |  |  |  |  |
| 3 weeks | 1800±80* | 0.310±133* | 158±138* | 82±5* | 243±46 | Nl. |
| 6 weeks | 2528±60* | 0.394±178* | 171±70* | 65±9* | 449±53* | Nl. |
| 12 weeks | 2194±57* | 0.304±126* | 258±103* | 70±2* | 436±63* | Nl. |
| GAA KO M/F-150mg n = 3 |  |  |  |  |  |  |
| 3 weeks | 436±40 | 0.307±138* | 120±73* | 86±1* | 442±75* | Nl. |
| 6 weeks | 1248±55* | 0.452±147* | 323±57* | 81±3* | 432±49* | Nl. |
| 12 weeks | 2549±68* | 0.397±180* | 355±104* | 67±15* | 344±54* | Nl. |
| WT |  |  |  |  |  |  |
| M/F n = 12-15 | 3,200±1423 | 0.569±296 | 264±101 | 60±16 | 334±96 |  |

\*p≤0.05 when compared to pre-treatment data by T-test (1-tail, 2-equal variance)

##### C6. Steady-state GAA levels in treated tissues

We measured steady-state GAA/NAG from the mice in above. Interestingly and not unexpected since the tobrhGAA is given orally, the SI sections contained the highest ratio and higher levels than wild-type-129/C57 (**Table 6**)(**127-137**).

**Table 6.**
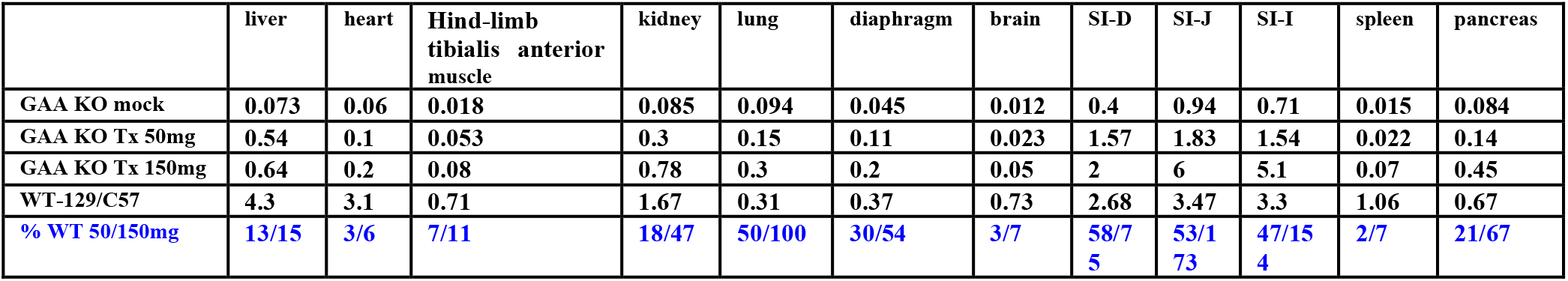

| TABLE 6 | liver | heart | Hind-limb tibialis anterior muscle | kidney | lung | diaphragm | brain | SI-D | SI-J | SI-I | spleen | pancreas |
| --- | --- | --- | --- | --- | --- | --- | --- | --- | --- | --- | --- | --- |
| GAA KO mock | 0.073 | 0.06 | 0.018 | 0.085 | 0.094 | 0.045 | 0.012 | 0.4 | 0.94 | 0.71 | 0.015 | 0.084 |
| GAA KO Tx 50mg | 0.54 | 0.1 | 0.053 | 0.3 | 0.15 | 0.11 | 0.023 | 1.57 | 1.83 | 1.54 | 0.022 | 0.14 |
| GAA KO Tx 150mg | 0.64 | 0.2 | 0.08 | 0.78 | 0.3 | 0.2 | 0.05 | 2 | 6 | 5.1 | 0.07 | 0.45 |
| WT-129/C57 | 4.3 | 3.1 | 0.71 | 1.67 | 0.31 | 0.37 | 0.73 | 2.68 | 3.47 | 3.3 | 1.06 | 0.67 |
| % WT 50/150mg | 13/15 | 3/6 | 7/11 | 18/47 | 50/100 | 30/54 | 3/7 | 58/75 | 53/73 | 47/154 | 2/7 | 21/67 |

##### C7. Pharmacokinetics and bio-distribution of Oral-ERT in GAA KO mice

Plasma and urine clearance rate with a 3x-dose (75 μg tobrhGAA) or whole seeds in GAA KO mice at 0, 1, 2, 4, 6, 8, 10, 12, 24 and 48 hr. TobrhGAA activity showed maximum serum GAA concentration (Cs) at 8 to 10 hr and peak urine excretion at 10 to 12 hr (**Fig. 4** and **5**). Bio-distribution (BD) in organs (skeletal muscle, heart, diaphragm, liver, spleen, brain, lung, sectioned SI and kidney) showed different levels and times of accumulation with showed greatest levels between 12, 24 and 48 hr (not shown)(**99,123,136-141**).

**Figure 4.**
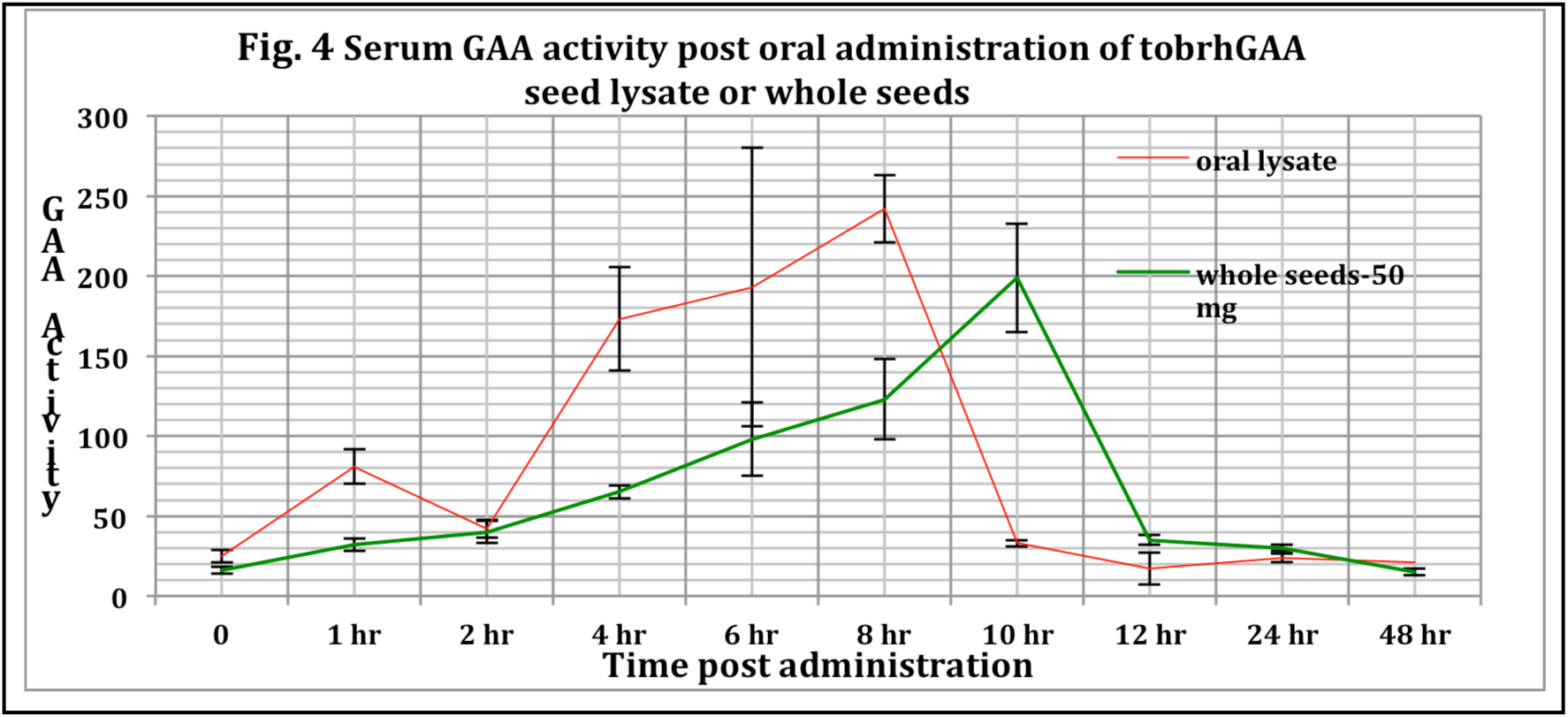
Pharmacokinetics in serum of Oral-ERT in GAA KO mice. Serum clearance rate with the 3x-dose (75 μg tobrhGAA) or whole seeds in GAA KO mice were collected at 0, 1, 2, 4, 6, 8, 10, 12, 24 and 48 hr. TobrhGAA activity showed maximum serum GAA concentration (Cs) at 8 to 10 hr.

**Figure 5.**
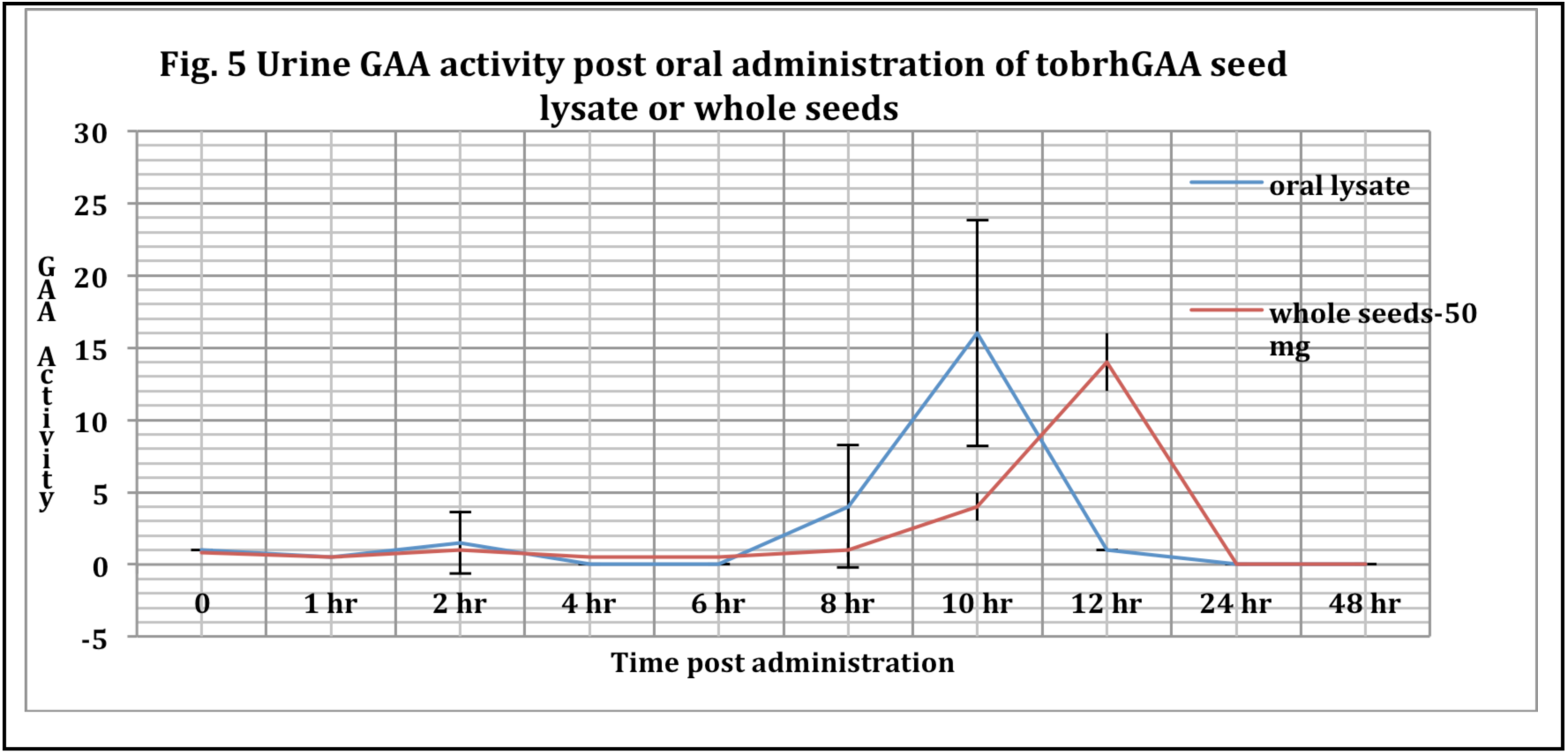
Pharmacokinetics in urine of Oral-ERT in GAA KO mice. Urine clearance rate with the 3x-dose (75 μg tobrhGAA) or whole seeds in GAA KO mice were collected at 0, 1, 2, 4, 6, 8, 10, 12, 24 and 48 hr. TobrhGAA activity showed maximum urine GAA excretion at 10 to 12 hr.

##### C8. Maximum tolerated dose

We determined the lethal dose in GAA KO mice (n = 4) starting at 10 mg-seeds/mouse, doubling daily to 320 mg. No weight loss or deaths were observed and CBCs remained unchanged.

## DISCUSSION

Currently, there is no effective treatment or cure for people living with PD. Sanofi-Genzyme, Inc. using a rhGAA (alglucosidase alfa) secreted from a CHO cell line has demonstrated moderate success in patients (**27**), however yearly costs are very high. Although alglucosidase alfa has been a wonderful first step in treating PD it has revealed subtle aspects that must be dealt with for successful treatment (**18,42-49**). ERT usually begins when the patients are symptomatic, however, all the secondary problems are already present which are further compounded by issues with alglucosidase alfa uptake in muscle (**Table 7**)(**18**). Lysosomal enzymes (such as GAA) are targeted to the lysosome by a mannose-6-phosphate recognition sequence that is exposed by posttranslational modification in the Golgi that may be the mechanism that extracellular GAA can be recycled and targeted back to the lysosomes. This mechanism will potentially allow rhGAA to be delivered to the cells or tissues and directed to the lysosome. However, some GAA may be taken up or recycled by endocytosis or a mannose-6-phosphate independent mechanism (**22-25**). In skeletal muscle there are a low abundance of the cation-independent mannose 6-phosphate receptors (CI-MPR) compounded by low blood flow plus low affinity of the CHO-produced alglucosidase alfa for the CI-MPR. Alglucosidase alfa is delivered to the lysosome by receptor-mediated endocytosis after binding of the M6P-glycan to the CI-MPR. Alglucosidase alfa has one M6P/enzyme (**142-144**) and 1,000-fold lower affinity (**145**). With alglucosidase alfa, the high level of infused enzyme is transient, leaving the patients with no therapeutic enzyme for the remaining 13 days. Thus, oral formulations of ground tobrhGAA seeds will allow patients to ingest daily in single or multiple doses conveniently at home to sustain a therapeutic dose of enzyme activity and to improve the quality of life.

**Table 7.**
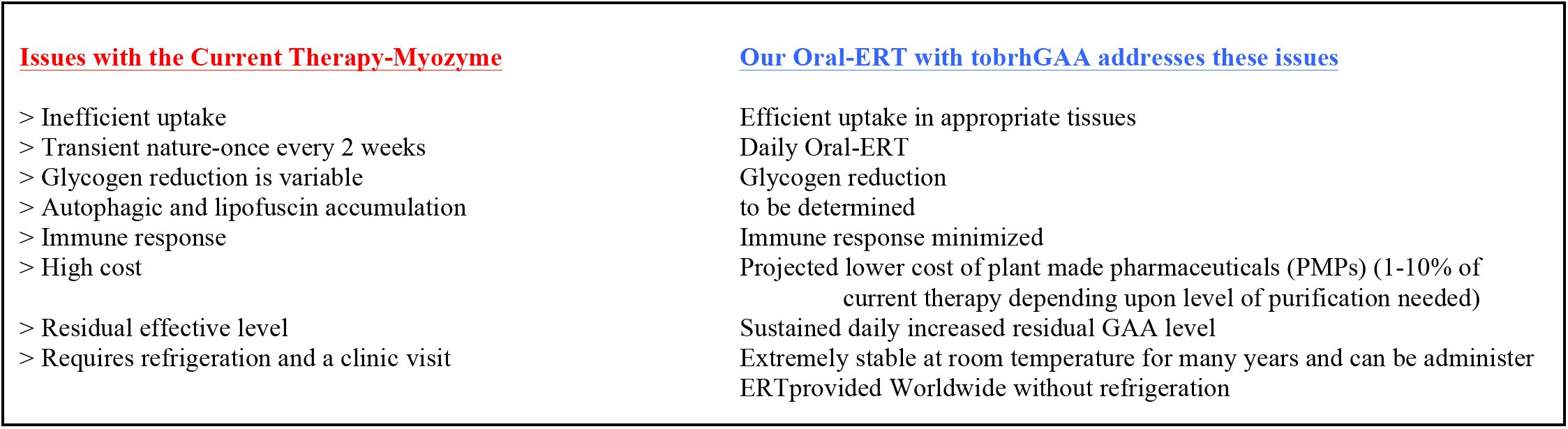

Investigators are studying the role of autophagy (**18,146-148**), an intracellular system for delivering portions of cytoplasm and damaged organelles to lysosomes for degradation/recycling in PD and the reduction of glycogen in skeletal muscle. A GAA and glycogen synthase 1 (*GYS1*) KO mouse exhibited a profound reduction of glycogen in the heart and skeletal muscles, a decrease in lysosomal autophagic build-up and correction of cardiomegaly. The abnormalities in glucose metabolism were corrected in the double-KO mice that demonstrate long-term elimination of muscle glycogen synthesis leads to a significant improvement of structural, metabolic and functional defects and a new perspective for treatment of PD (**17**). Lim et al. (**148**) reactivated mTOR in PD mice by TSC knockdown resulted in the reversal of muscle atrophy, removal of autophagic buildup plus aberrant mTOR signaling can be reversed by arginine alone (**147-149**). Jung et al. (**150,151**) produced and characterized a rhGAA in a transgenic rice cell suspension culture with high-mannose glycans and was similar to the CHO derived hGAA. Sariyatun et al. (**152**) produced a hGAA with a paucimannose structure in *Arabidopsis alg3* cell culture for which M3 or mannose receptor-mediated delivery.

Seeds may be a better vehicle for Oral-ERT for lysosomal diseases, such as PD, compared to intravenous systemic delivery. Seeds and not other plant tissues and organelles, contain the metabolic machinery necessary for correct glycosylation, processing, phosphorylation and synthesis of complex enzymes and proteins, enzymes or proteins plus are protected or shielded from digestion in the stomach and small intestine and can be administered daily in single and multiple doses and seeds provide long-term stable storage of recombinant enzymes. A number of biotechnology companies have tried to mass produce a rhGAA from different platforms. A consensus at a 2019 US Acid Maltase Deficiency conference at San Antonio, TX, suggested that a multi-pronged approach including gene therapy, diet, exercise, etc. must be evaluated for a successful treatment of PD. Prompted by studies of oral formulations of edible tissues in broccoli sprouts, corn/maize, pea, rice, tobacco and tomato for pharmaceutical applications including vaccination to induce effective mucosal immune tolerance and immune reactions, GI, bacterial/viral infections, allergies, asthma, diabetes, endocrine-associated diseases, hypersensitivity, elevated BP, cholera, leucopenia, cancer and RA (**120,153-172**), we hypothesized that rhGAA from edible plant tissues offers an cost-effective, innovative and safe strategy for Oral-ERT for PD. We investigated the potential of genetically engineered “edible plant tissues” as an alternative large-scale production system that overcomes the high cost of producing rhGAA in CHO cells. Oral-ERT can be safer than infusion and be ingested in a pill/capsule format at frequent intervals daily to maintain a therapeutic dose of enzyme to achieve long-term clinical efficacy. This protocol will be the second oral ERT plant-made pharmaceutical (PMPs)(**74,89-91,96**). Thus, to provide a less expensive alternative, we generated a rhGAA produced in tobacco seeds for ERT of PD. We found that the tobrhGAA compared favorably and was superior on many aspects to alglucosidase alfa (**Table 7**). After scaling-up, we estimate that the yearly cost for an adults living with PD would be $3,000 or <1% of the cost of alglucosidase alfa which ranges from $250,000 to $650,000 per adult patient per year depending upon weight. Tisdale et al. (**173**) in a retrospective study of medical and insurance records indicates health care costs for people with a rare disease have been underestimated and are 3 to 5x greater than the costs for people without a rare disease. Only about 10% of rare diseases have an FDA-approved therapy with most of the 7,000 to 10,000 affect children, adolescents and young adults. Individually rare diseases affect only a few hundred to a few thousand worldwide, however, collectively affect an estimated 25 to 30 million people in the USA. The cost per patient per year (PPPY) with a rare disease exceeded costs for non-rare diseases patients. PPPY costs ranged from $8,812 to $140,044 for rare diseases patients compared to $5,862 for those without a rare disease. Extrapolating the average costs for the 25 to 30 million individuals with rare diseases in the USA results in a total yearly direct medical costs of approximately $400 billion, which is similar to annual direct medical costs for cancer, heart failure and Alzheimer’s disease.

Successful demonstration of Oral-ERT with tobrhGAA will have significant clinical applications and shift current clinical practice paradigm by offering a safe, stable at room temperature and affordable lifelong Oral-ERT for PD in the USA and World-wide especially in areas where access to a clinical setting for biweekly IV administration of Myozyme is lacking. We estimate that an adults living with PD patient will need to take 3-4 grams daily at a cost of 0.1% to <1% of Myozyme depending on how it is branded and marketed to maintain a sustained GAA level of 5-10% of normal.

## Abbreviations

AMD: acid maltase deficiency
rhGAA: recombinant GAA
tobrhGAA: recombinant GAA produced in tobacco seeds
GAA: acid maltase
ERT: enzyme replacement therapy
Oral-ERT: oral enzyme replacement therapy

## ACKNOWLEDGEMENTS

None

## Legends

**Graph 1.**
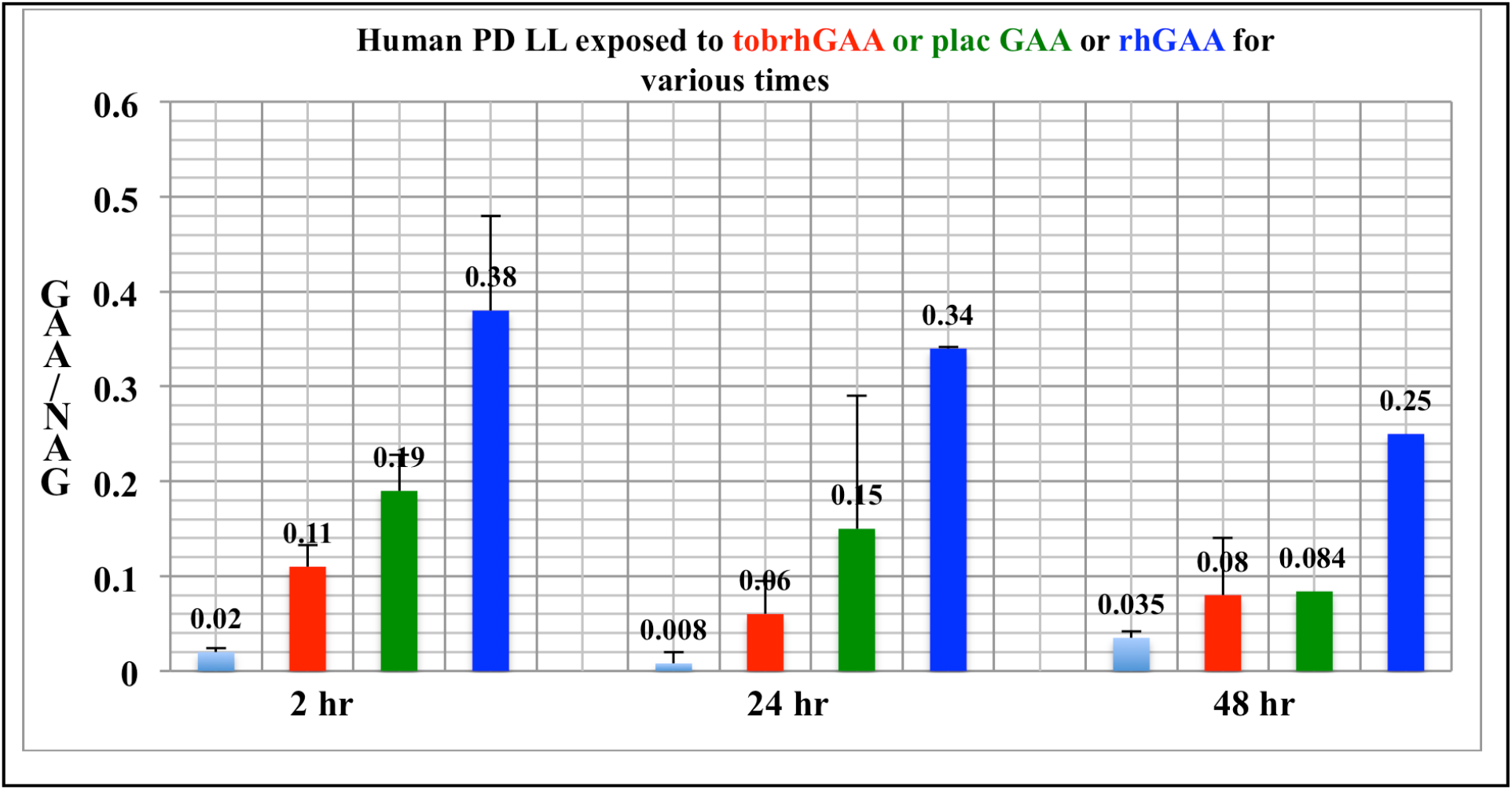
Uptake of tobrhGAA in human PD lymphoid cell lines. Human lymphoid cell lines from infantile or adult PD (GM6314, GM13793, GM14450) were exposed to equivalent amounts of tobrhGAA, placental GAA or a rhGAA. Cells were harvested after 2, 24 and 48 hr and assayed for GAA and NAG. All treatments were significant at p = ≤0.05.

**Graph 2.**
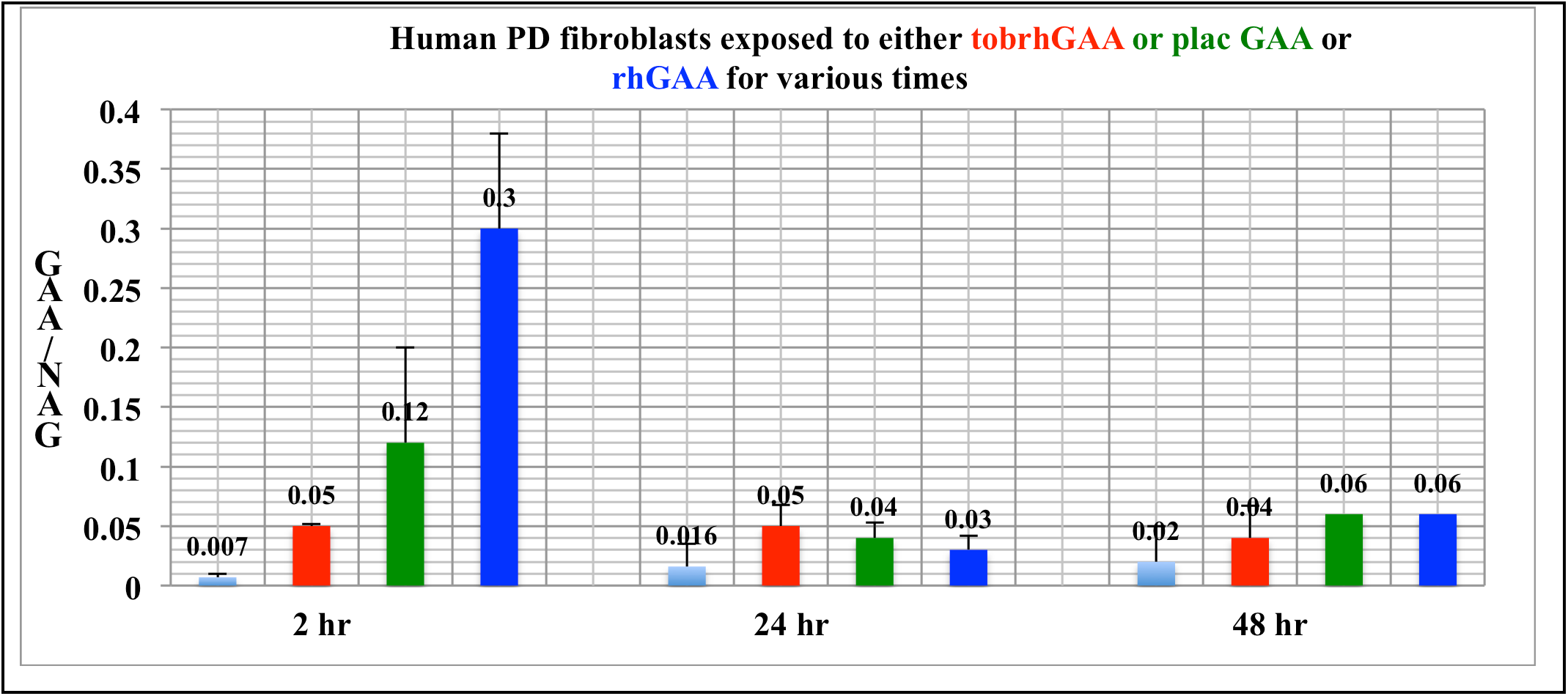
Uptake of tobrhGAA in human PD fibroblast cell lines. Human fibroblast (GM4912, GM1935, GM3329) cell lines from infantile or adult PD were exposed to equivalent amounts of tobrhGAA, placental GAA or a rhGAA. Cells were harvested after 2, 24 and 48 hr and assayed for GAA and NAG. All treatments were significant at p = ≤0.05.

**Graph 3.**
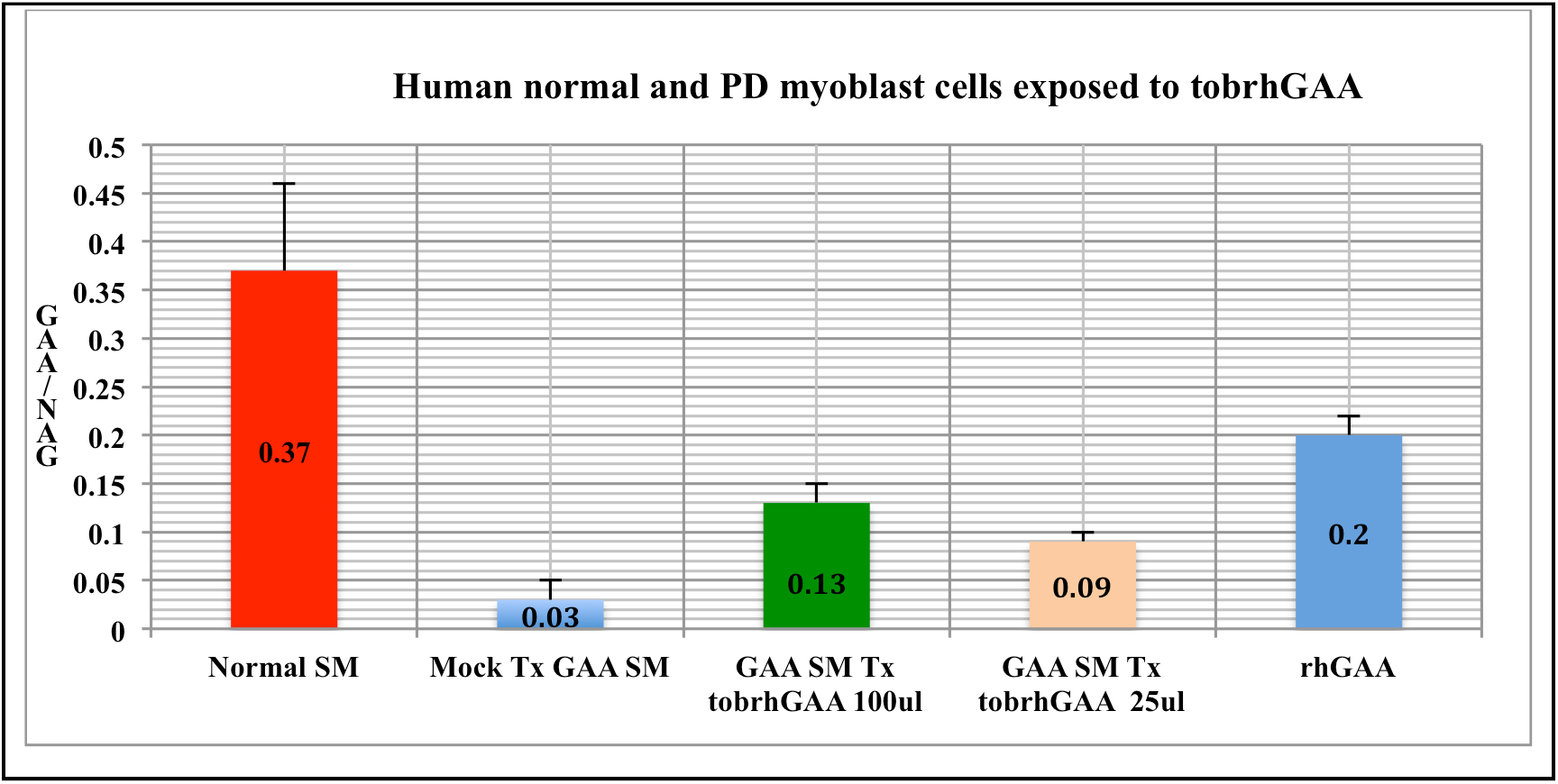
Uptake of tobrhGAA in a human PD myoblast cell line. A human PD myoblast cell line was exposed to equivalent amounts of a lysate of tobrhGAA or an rhGAA for 48 hr and assayed for GAA and NAG. We found that tobrhGAA increased GAA to 24-35% of normal (mean±SD).

**Graph 4.**
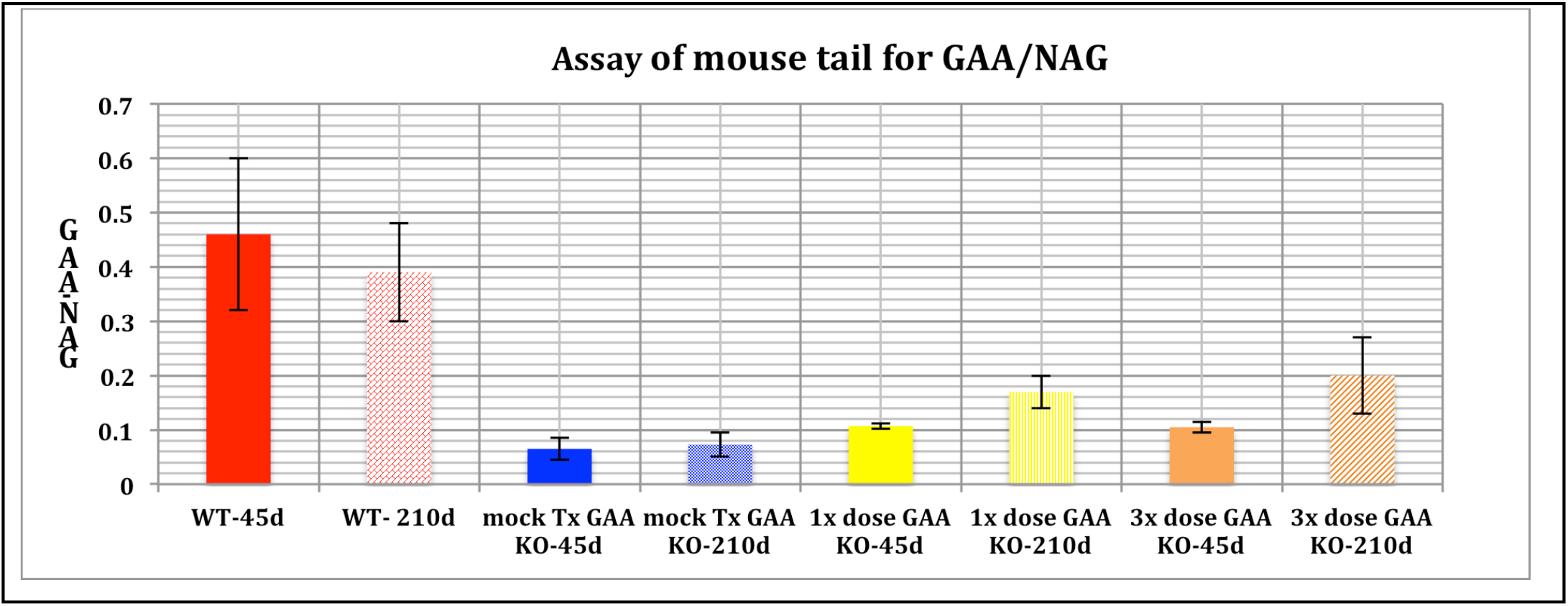
Taken up by a distant tissue. We determined if the tobrhGAA was taken up by a distant tissue-tail without sacrificing the mice and assayed for GAA at 45 and 203 days. Wild-type mice had GAA activity of 0.46±0.14 (SD), mock treated GAA KO had 0.064±0.021 and treated GAA KO had 0.10 to 0.2±0.05 for 1x or 3x dose (p = ≤0.006 treated vs mock treated).

**Graph 5.**
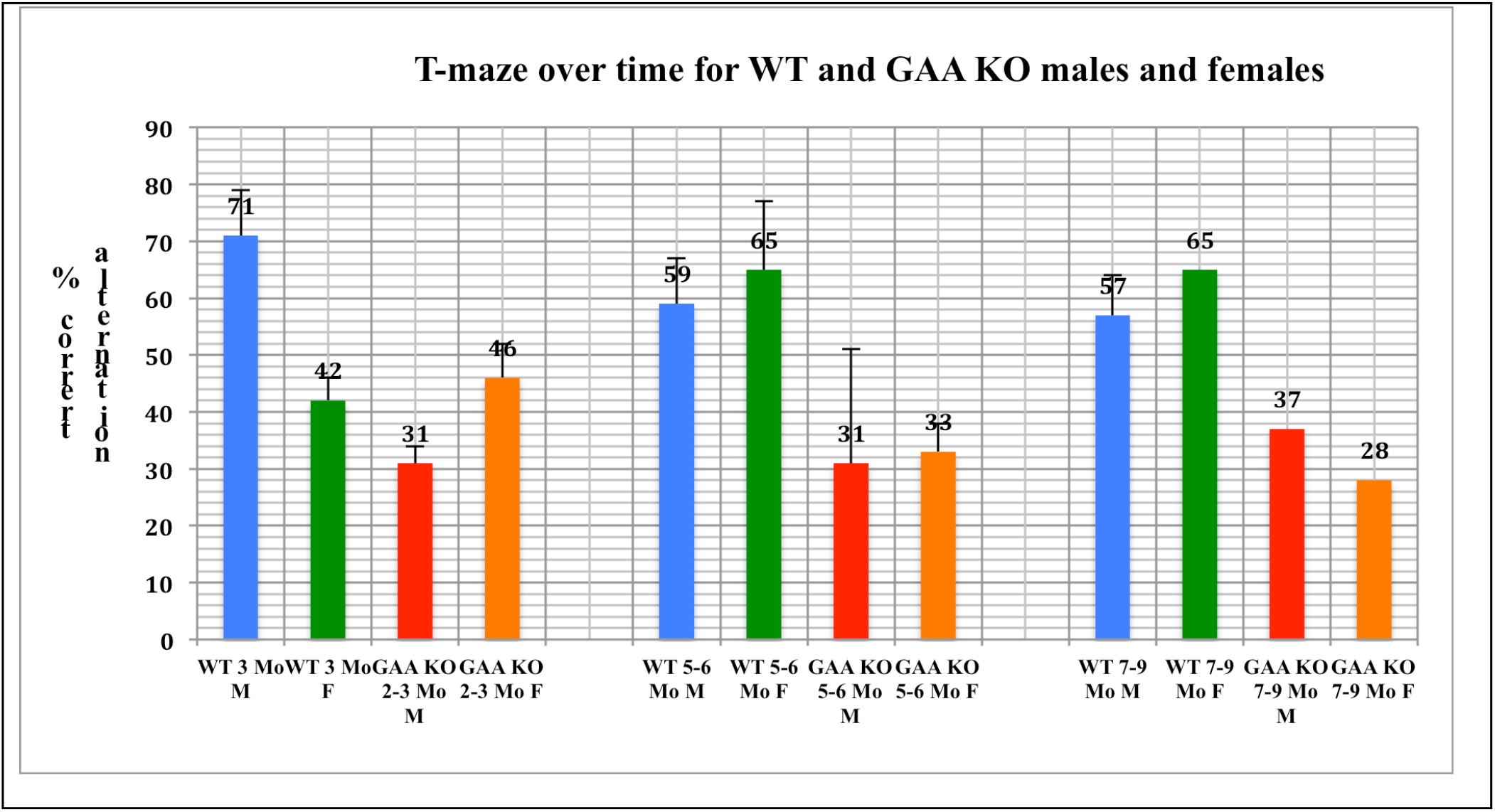
Assessment of spontaneous alternation in wild-type and GAA KO mice. We assessed spontaneous alternative learning for cognitive ability in the T-maze in both male and female GAA KO mice and WT-129/C57 mice from 2-9 months of age. We found that deficiency in spontaneous learning appeared by 2-3 months in male and 3-4 months in female GAA KO mice. All conditions were significant (p = ≤0.05).

